# Accurate Isothermal Amplification Reaction Rate Determination Using High-Frequency Sampling

**DOI:** 10.1101/017160

**Authors:** Thomas Beals

## Abstract

**Background:** Nucleic acids quantification by amplification is currently done primarily by real-time amplification for relative quantification, or by statistical inference from replicated endpoint assays for absolute quantification. The polymerase chain reaction (PCR) has been the dominant amplification technology, although alternative isothermal technologies have been described. Theoretical analysis of amplification kinetics and of amplification data interpretation have almost exclusively considered the PCR.

**Results:** Real-time measurements of isothermal amplification reactions can be made continuously, in contrast to the discrete per-cycle measurements of real-time PCR. Isothermal ramified rolling circle amplification (RAM) reactions were measured at frequent intervals, and amplification data subsets were fitted to an exponential amplification model. Signal-change-over-time slopes and time-zero signal intercepts were derived from the chosen subset data. Slope measurements were sufficient to determine a reaction rate (the isothermal equivalent of PCR efficiency) for each reaction. Analysis of slope and intercept together suggest that amplification reactions that were initiated from a single target molecule can be distinguished from reactions that that were initiated from greater than one target molecule.

**Conclusions:** The constant reaction environment of isothermal nucleic acid amplification allows continuous monitoring of reaction rate. Functional regions of interest in real-time data can be determined directly from the data. Accurate per-reaction efficiency can be readily measured. Improved estimation of low target copy number should improve quantification efficiency.

## Introduction

Nucleic acid amplification can provide sensitive and specific molecular detection and quantification. Since its introduction PCR has been the dominant amplification technology; its widespread use has stimulated improvements in enzymes, instrumentation, and analytical methods. Initial use of PCR was largely qualitative: positive target detection, or inference of target absence above a detection limit. PCR was made more quantitative with the advent of real-time PCR, notably for relative quantification that estimates a target concentration ratio between two samples. More recently, digital PCR (dPCR) has allowed absolute quantification – the estimation of the number of target molecules in a given sample volume.

Target quantification by real-time PCR has motivated a substantial literature (1, 2; online bibliography 3) regarding the determination of PCR efficiency the amount of target amplification per PCR cycle. PCR efficiency determination is problematic because it must be estimated from a small number of data points. Digital PCR typically is not performed in a real-time instrument; dPCR interpretation is based on analysis of ratios of amplification reactions that contain no target molecules, to reactions that contain one or more target molecules.

A variety of non-PCR nucleic acid amplification methods have been described (4). Isothermal amplification methods have in common a constant – that is, non-thermo-cyclic – reaction environment, and isothermal real-time reaction signals can be monitored continuously. A constant reaction environment allows the collection of sufficient observations for an amplification rate to be determined more precisely than is currently possible for PCR. Results from isothermal amplifications that were monitored in real time, including target concentration ranges used in dPCR, are presented here. Statistics derived from replicate reaction data are interpreted as allowing the discrimination of reactions that amplified a single target molecule, from reactions that were initiated from two or more molecules.

## Results

The RAM reaction mechanism, like the PCR reaction mechanism, predicts exponential amplification product increase over time (5). Figure 1 illustrates real-time monitoring of amplification reactions. The idealized amplification reaction is assumed to be exponential from the start, although the signal from early reaction products is less than the baseline signal – the noise threshold of the detection system (Figure 1; A,B). The interval where the observed signal changes dynamically with respect to time is initiated when accumulated amplification product produces a signal that exceeds the background level. Exponential signal change ends as the dynamic signal transitions to a static plateau; that transition may be caused by one or more factors such as reagent depletion below optimum concentration, loss of enzyme activity, or reactant accumulation that causes a transition to a less favorable reaction environment. An assumption common to PCR analysis and to the analysis method presented here is that there exists a region, or subset, of observable data from which the unobserved reaction course (Figure 1B) can be inferred.

**Figure 1.**
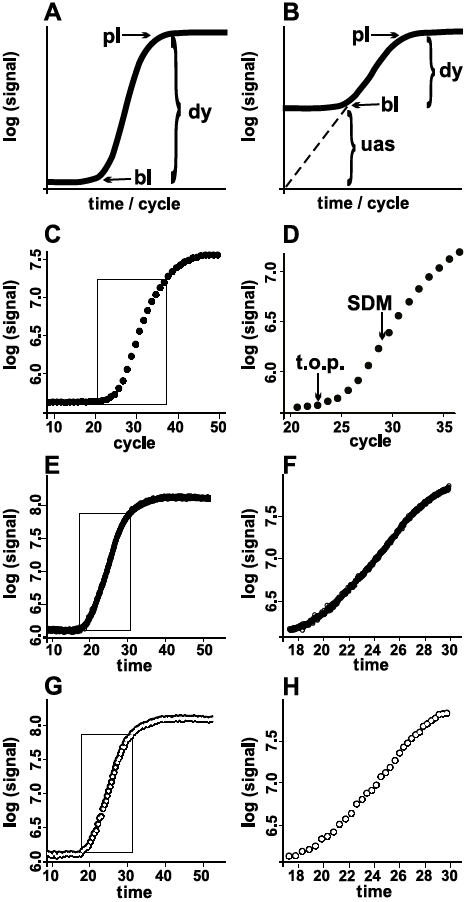
Real time exponential amplification: examples and interpretation. A) Conceptual form of a real-time amplification data. The baseline (bl) is due to tracer fluor or instrument noise. The signal changes significantly with respect to time in the dynamic (dy) region; the signal does not change significantly at the plateau (pl). B) The dashed line indicates an unobserved assumed signal (uas). The reaction becomes observable when it generates a signal that is greater than the baseline signal. C, D) Data from a PCR. The single data point obtained per cycle is represented by a solid circle. An inset rectangle in Figure 1C indicates an area that is shown in Figure 1D. Takeoff point (t.o.p) and second derivative maximum (SDM) are indicated by arrows. E, F, G, H) Data from a RAM reaction. Open circles represent data points; individual open circles are not distinguishable in Figures 1E, 1F. Inset rectangles in Figures 1E, 1G indicate areas shown in Figures 1F, 1H. Figures 1G, 1H are drawn from the same data as Figures 1E, 1F but allow visualization of individual data by display of one out of every twenty data points.

Figure 1 shows that real-time RAM and PCR reactions produce similar sigmoidal signal vs. time forms; each having a baseline, a dynamic region of positive-slope signal change over time, and a plateau. However, the PCR signal is measured (or summarized) once per cycle, while the isothermal RAM reaction signal can be measured as close to continuously as instrumentation allows – here, each isothermal RAM reaction has been measured at 0.5 to 3-second intervals. The data density obtained by frequent data collection during isothermal amplification, by contrast to the PCR signal, is illustrated in Figure 1 (C H). Data from a PCR reaction is shown on Figure 1 C, and that reaction’s dynamic region (from transition out of baseline to initial slope-decrease) is shown on Figure 1D. Numerically, the data point ratio of Figure 1F:Figure 1D is about 38:1. The region of PCR exponential signal change on Figure 1D is marked by a ‘takeoff point’ (6), to a second derivative maximum (7). Inference of informative statistics from the PCR’s single data point per cycle remains an analytical challenge (7, 8).

Figures 1E, 1F show a RAM reaction and a detail section that is comparable to Figures 1C, 1D. Although the plotting character is specified as an open circle, the data density is such that the plot appears as a solid line. Figures 1G, 1H are drawn from the same data, but only one in twenty data points is displayed, to show individual data points. Data analysis was done by a first pass through each reaction’s data to provided an estimate of the amplitude of the dynamic region; that estimate was used to partition the data into a no-amplification set and a positive-amplification set. A second pass analysis for positive amplifications is illustrated using the data shown in Figure1, panels E-H.

PCR data analysis methods that derive reaction rate estimates were not used for these data because those PCR methods were not designed to take advantage of these larger data sets. However, as for PCR, analysis of these data required choosing a data subset from the dynamic region as representative of the reaction rate. Representative data subsets were chosen by first finding a maximum first derivative of log-linear signal vs. time, using numerical differentiation with parameters that smoothed the local data noise. A log-linear model was then fit to a fixed interval around the first derivative maximum. This initial data evaluation method was simple to implement and provided objective statistics for comparative evaluation of replicated amplification reactions.

For each amplification reaction, the outcome of the analysis process is a collection of statistical parameters derived from a data subset. These parameters include the slope and intercept of a line fitted to the data points within the subset, the endpoints of the fitted region, and the signal variance within the fitted region. Figure 2A shows a 61 data-point segment that was identified algorithmically from the data shown in Figure1E-H, as well as a line that was determined by the slope and intercept of the log-linear model. Figure 2B shows the dynamic region of the amplification data, with an extrapolation of the fitted line, and a graphic indication of the position of the subset data.

**Figure 2.**
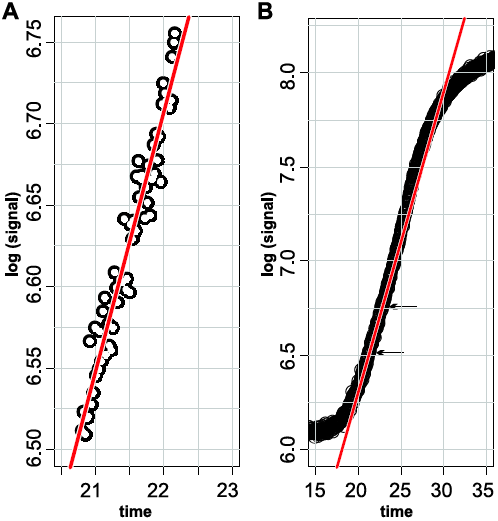
A best-fit line determined from the RAM reaction data shown in Figure 1. A) Open circles represent data-points within an algorithmically selected data subset. The red line was determined by a log-linear fit of signal vs. time within the selected subset. B) The line determined by the data shown in Figure 2A is extrapolated within a plot of the dynamic region data. Arrows mark the bounds of the data subset shown on panel A.

To assess the utility of these analysis methods for quantification, reagents were prepared to allow comparison of physically measured target concentrations with concentration estimates from amplification reactions. A stock of single-stranded DNA (ssDNA) RAM-target circles that was substantially free of linear DNA precursors was prepared (Figure 3); spectrophotometric measurements of this material provided an estimate of 2.5 × 10^11^ ssDNA targets per microliter. The target stock was diluted, with each dilution step monitored by replicated weighings on an analytical balance, so that the volumes as measured by pipetting could be refined by adjusting for the measured liquid mass.

**Figure 3.**
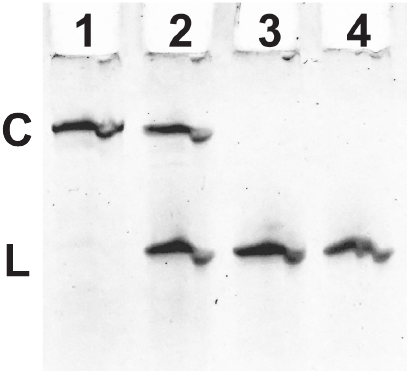
Acrylamide gel image of single-stranded DNA circles that are substantially free of linear precursor. 1, single-stranded circular DNA after exonuclease treatment and purification; 2: same starting material and processing, but mock-exonuclease treatment; 3, 2 pmole linear DNA precursor; 4, 1 pmole linear DNA precursor. C: single-stranded circular DNA; L, linear DNA.

Replicate amplification reactions were performed on five levels of sample dilution, from a nominal 128 target molecules per reaction to a nominal 0.5 target molecules per reaction, with 4-fold dilutions between each level. Table 1 shows the number of reactions per dilution level, the nominal target concentration per reaction, and the amplification-failure ratios for the two lowest concentrations. (As for dPCR, failure means no amplification; and that failure is interpreted as the absence of any target molecules.) The number of replicate reactions per dilution level was chosen based on prior experience with the system; lower replicate numbers were chosen for higher target concentrations where lower variance was expected. Table 1 shows the Poisson means (calculated as for dPCR analysis) for the two dilutions for which there was an appropriate proportion of positive-amplification and non-amplified reactions. A nominal 2 target molecules per reaction dilution was estimated to have ∼1.2 target molecule per reaction, and a 0.5 nominal target molecules per reaction was estimated to have ∼0.65 target molecules per reaction. Multiplication by the final calculated dilution factor calculated from the two dilutions yields target stock concentration estimates of ∼2.05 × 10^11^ target molecules per microliter to 4.60 × 10^11^ target molecules per microliter.

**Table 1.** Table 1. RAM amplifications; experiment design and outcomes.

| Nominal count | 128 | 32 | 8 | 2 | 0.5 | 0 |
| --- | --- | --- | --- | --- | --- | --- |
| Replicates | 8 | 8 | 12 | 16 | 44 | 8 |
| Number fail | 0 | 0 | 0 | 4 | 19 | 8 |
| QC fail | 0 | 0 | 0 | 1 | 4 | 0 |
| Fraction fail | 0 | 0 | 0 | 5/16 | 23/44 | 1 |
| Poisson mean | n/a | n/a | n/a | 1.16 | 0.65 | n/a |

To determine whether analysis of real-time data could add interpretive value to conventional Poisson failure analysis, the analytical methods described above were applied to each positive amplification. Figure 4 shows sigmoid amplification signals (as in Figure 1), and lines determined by log-linear fitting to defined data subsets (as shown in Figure 2) for each signal (Figure 4, left panels). The right panels of Figure 4 show extrapolated log-linear fit lines without the amplification signal. Published studies (9) have scored and compared realt-ime data such as the signal traces shown on Figure 4 by using response time (Rt) – a measure that, like the Ct of PCR, is the time when a signal-trace dynamic region crosses a given signal level (Figure 5A). Quality control (Table 1, “QC fail”) are reactions with response times over 40 minutes. Statistical analysis augments a previous (9) quality control criterion of Rt greater than 40 minutes, but is not further analyzed here.

**Figure 4.**
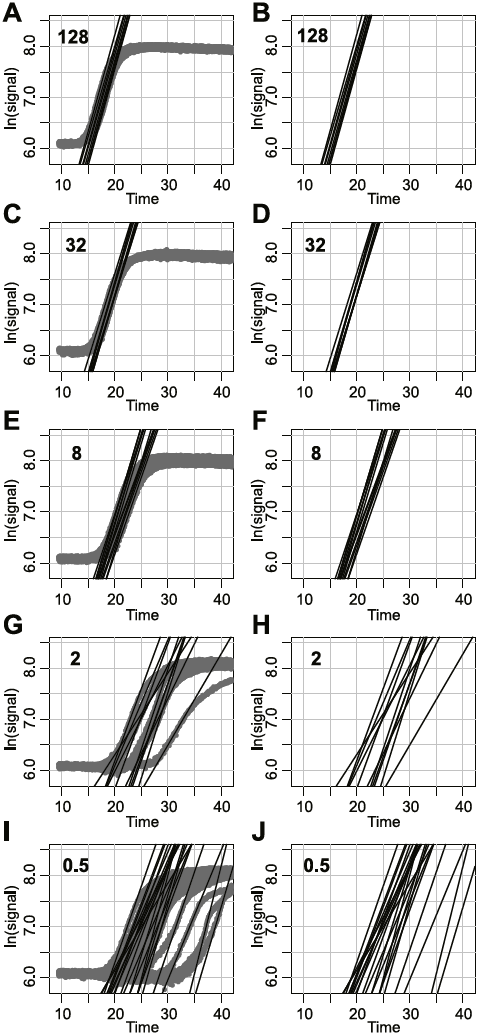
RAM real-time data and extrapolated fitted lines. Left panels A, C, E, G, I show sigmoid-form real-time trace data. A fitted line through the dynamic region of each real-time data trace is shown. The extrapolated lines are shown without the sigmoid trace data on right-side panels B, D, F, H, J. The nominal template numbers per reaction and per panel are Panels A, B: 128; C,D: 32; E, F: 8; G, H: 2; I, J: 0.5.

**Figure 5.**
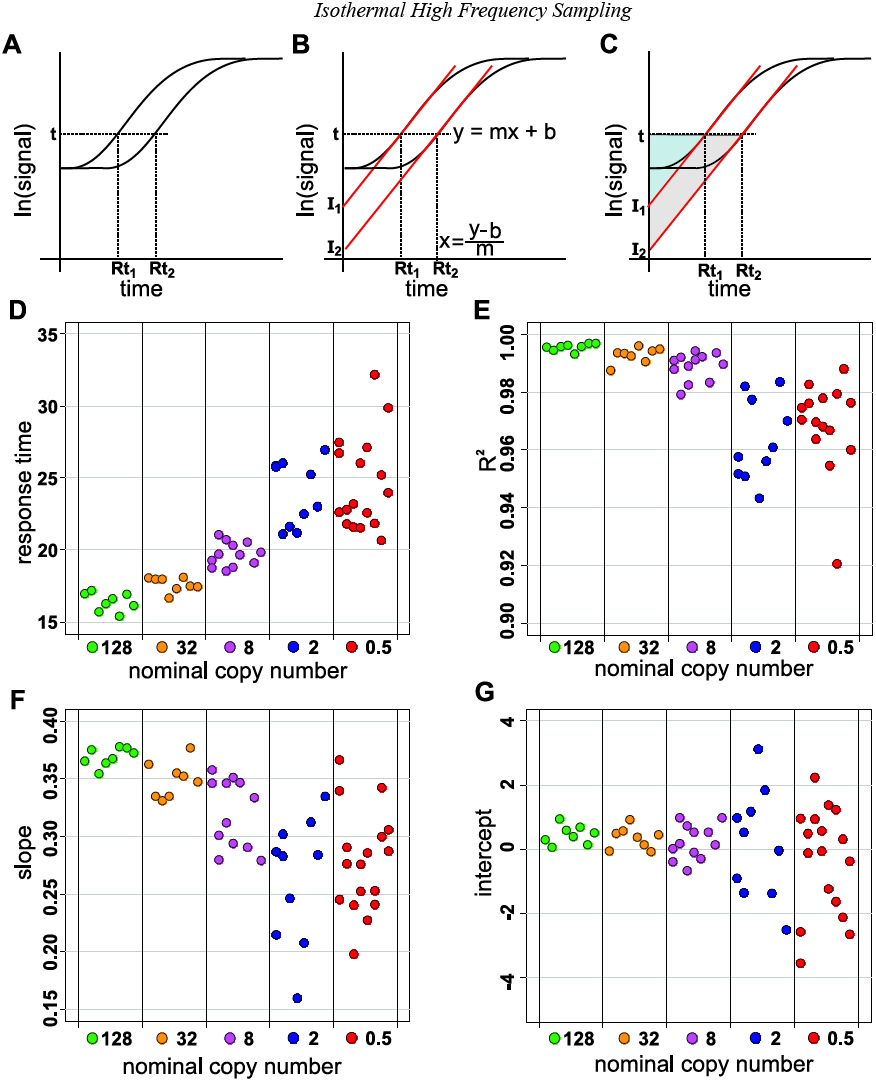
Components of response-time. (A-C) Two idealized signal-traces with red lines indicating fitted linear models. A. Response time (Rt) is the time of the intersection of a signal level (t) and a signal trace. B. Fitted lines are specified by a conventional linear formula y*(signal)* = m*(slope)* * x*(time)* + b*(y-intercept)*. C. Similar triangles are defined by fitted lines with equal-slopes, the fitted line’s intersection with the signal level (t), and the vertical axis. (D-G). Amplification data analysis. D. calculated response times for each of 5 nominal target levels. E. Correlation coefficient R2 for fitted lines. F. Slope component of response-times. G. Intercept component of response-times.

Response time as defined does not account for possible differences in slope, and Rt comparisons that assume equal slope could be misleading. The slope and intercept parameters can be used to determine a response time algebraically for any chosen signal level (Figure 5B), once a log-linear model has been determined. For equal-slope models, the time-zero intercept for comparison of two reactions is an alternative to the Rt. Figure 5C shows that fitted lines with equal slopes form similar triangles with proportional sides. This mathematical relation implies that a change in response time will have a proportional change in intercept. The time-zero intercept has the physical interpretation of being the predicted signal for that reaction’s input targets before amplification – that is, a measure of the initial target number; the goal of the assay (as recognized for PCR; 10).

The dynamic regions of real-time traces shown in the left panels of Figure 4 are generally earlier – have lower response-times – for greater target numbers. Figures 5D – 5G and Figure 6 show response times and statistical components from fitted models to further explore response time dependence on target number. Figure 5D shows response time as a function of nominal copy number; response time is both later and more variable for lower nominal copy number reactions. While the correlation coefficient of the fitted lines may make some contribution to response-time variance, Figure 5E shows that although R^2^ for the log-linear models decreases with lower nominal copy number, even at the lowest (nominal 0.5 targets per reaction) level, most model fits have R^2^ greater than 0.95.

**Figure 6.**
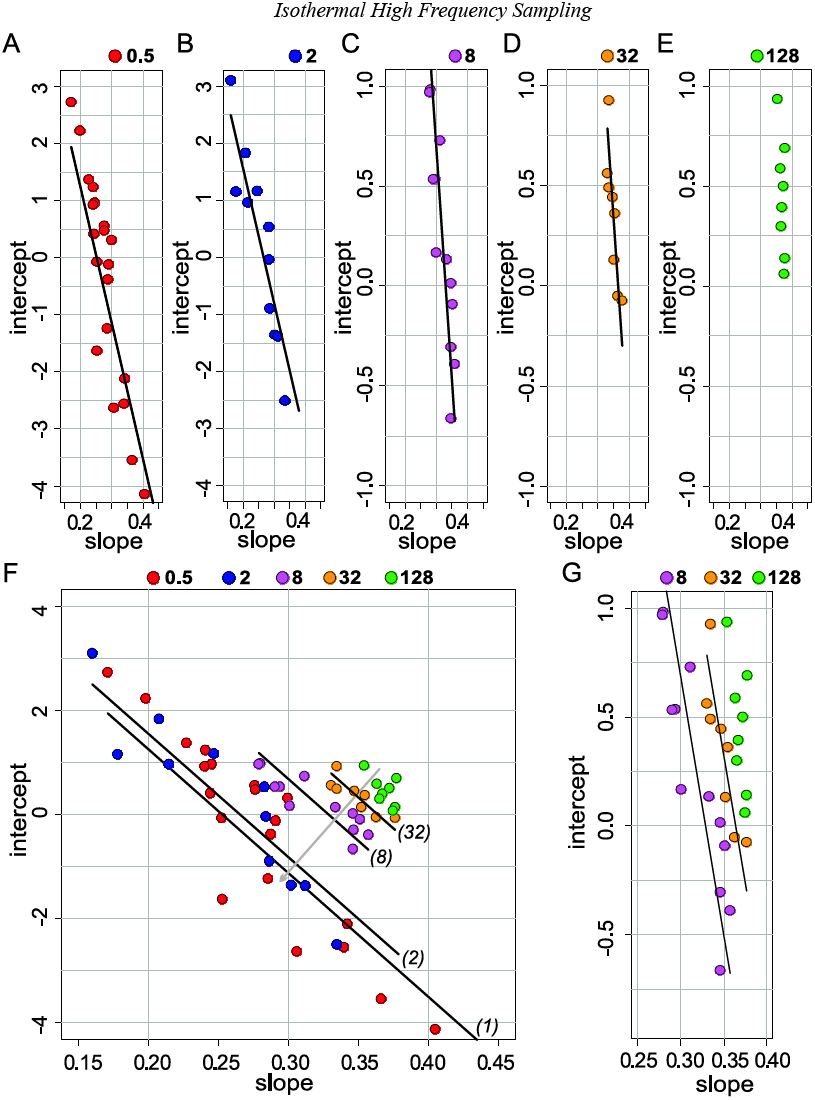
Slope-intercept plots from RAM ttarget dilutions. Slope and intercept data from RAM reactions on five nominal template levels are shown as color-coded points. A single linear equation with slope and nominal template number as factors was used to generrate fitted lines. Nominal template levels per reaction are: A, 0.5; B: 2; C, 8; D, 32; E, 128. Panel F is a composite of the separate data from panels A through E; fitted lines are labeled with their copy number model argument. A grey line perpendicular to the fitted lines indicates a copy-number gradient. The vertical scales of panels C, D, and E are identical, and panel G is a composite of the data from panels C, D, and E.

Figures 5F and 5G show the slope and intercept components of the fitted models. Figure 5F shows an apparent dependence of slope (reaction rate, for isothermal reactions) on nominal copy number; the reaction rate decreases as nominal copy number decreases. A decrease in slope, all else being equal, will increase the y-intercept. Figure 5G shows increased variance in y-intercepts with copy number, but there is no obvious functional dependence on y-intercept with copy number.

A slope decrease with copy number appeared to be consistent with a proposed mechanism for the RAM reaction (discussed below, and (5)), but the apparent non-correlation of intercept and copy number was unexpected: as noted above, for equal slope reactions, y-intercept is expected to be proportional to copy number; and Figure 5D showed that response time increased with decreasing copy number. To visualize possible relations between slopes and intercepts of these fitted models, Figure 6 plots slopes vs. intercepts; the intercepts on a vertical axis, and the slopes on a horizontal axis. Results from the five target levels are plotted separately on Figure 6 A-E, and together on Figure 6 F. The three highest concentrations are plotted together on Figure 6G.

Ideal amplifcation reactions with a constant reaction rate that was independent of initial copy number, plotted as on Figure 6, would distribute intercepts vertically at the constant slope. Figure 6E visually approaches that ideal; Figure 6G combines plots of the three highest concentrations. The slope change as a function of copy number can be seen as the tendency of data points to shift rightward as the copy-number increases (Figure6 A-E and G); as well, the intercept variance decreases with increasing copy number.

Figure 6F combines Figures 6A-E. The scale of the horizontal ‘slope’ axis is expanded, but shows the same range as the upper panels. Lines superimposed on the data points were generated by a statistical model of intercept as a function of slope and of nominal target number per reaction; the slope-intercept data points cluster around diagonals that can be visualized by the fitted lines. The terms of the equation that generated the fitted lines shows the influence both of nominal target number and of slope as determined for each reaction. The diagonal patterns show that the 0.5 and 2 nominal-copy-number data points assort along those diagonals in numbers that are consistent with Poisson expectations for the calculated input copy-number. Figure 6F is interpreted as showing that copy number is roughly correlated with distance along a line that is perpendicular to the fitted diagonals (Figure 6F, light grey line). Since the two lowest target reactions included some non-amplified wells interpreted as no-target reactions, a substantial proportion of the 0.5 nominal copy number reactions are expected to have been amplified from a single target moleclue.

## Discussion

The primary goal of this communication is to demonstrate the utility of continuous, or high-frequency, sampling of isothermal amplification reaction signals. That utility is shown here with isothermal RAM reactions, but high-frequency sampling of any isothermal reaction should be feasible if that reaction can be monitored in real time. The value added by high-frequency sampling is the creation of a larger reaction data sample size with concomitant gains in statistical significance. Figure 1 contrasts the data density difference between PCR and the isothermal method used here; Figure 1 G, H shows one out of twenty data points to allow visualization of individual data points, and to illustrate the data density obtained in contrast to PCR. The distinction between PCR data collection and high-frequency isothermal sampling is that PCR amplification is measured at discrete integer cycle numbers, whereas isothermal reaction rate measurement can be done continuously – as if the fractional (non-integer) cycles reported in PCR Ct or Cq results could be observed directly.

Data collection as described should not require changes in sample preparation, reaction chemistry, reaction time, or instrumentation hardware (although software versatility will be required); after a real-time instrument is set to record amplification data at an appropriate frequency, workflow should proceed unchanged. For any quantitative amplification procedure, however, the quality of the result will depend on accuracy in dilution, and quantitative isothermal amplifications may benefit from automated procedures that have been developed for digital PCR sample preparation. The data shown here were collected in an instrument designed for PCR; instrumentation designed for isothermal high-frequency sampling (IHFS) may provide higher quality data by decreasing measurement variance and by optimizing sampling frequency.

The PCR literature has been invaluable for conceptual interpretation of IHFS data, but the computational tools described for PCR did not appear to be applicable to the analysis of IHFS data. In broad overview, analogous tasks to a PCR analysis were performed: data collection, individual reaction analysis, and interpretation of results in light of experiment design. Data collection differs from previously described methods as appropriate to reaction specifics and instrumentation settings. The analysis method described here is conceptually simpler than many PCR analyses: a data subset that is likely to represent the reaction is chosen, and descriptive statistics are derived from that segment. A fixed-size segment that was determined by the first derivative maximum was used for this analysis, after exploration of several alternative methods for selection of representative data subsets. While almost certainly not an optimal method, the chosen method was simple and gave interpretable results. Figure 2 shows an example data subset, a fitted line, and the data-point density around the fit. Statistically, and perhaps intuitively, there appears to be enough data for good fit predictions. The data density and linearity makes the analytical computation very quick. The analysis process output is a per-reaction data table that includes slope, intercept, measures of goodness of fit, and the bounds of the fitted segment; that data table was the basis for the interpretation of the experiment.

Figure 4 shows the observed real time data – a log-transformed graphical representation with extrapolated fitted lines is shown for each reaction. As the nominal input target number per reaction decreases, the rightward shift of the signals’ dynamic region is apparent, as well as the greater signal variance. The signal change with respect to target level seen on Figure 4 is quantified by the response time as defined on Figure 5A, and as shown on Figure 5D. A response time can be calculated from the slope and intercept parameters of the fitted line; the algebraic transformation that allows the calculation of a response time given a signal level is shown on Figure 5B. The proportionality of Rt values and corresponding intercept values for equal-slope fitted lines is made with a geometric illustration on Figure 5C. The interpretation of the time zero intercept as the predicted signal level of the unamplified target was made by Rutledge and Côté (10); however as Tellinghuisen and Spiess note (7) the intercept is a substantial extrapolation, whereas the intersection of a signal trace with a given signal level is a point that can be directly identified. That note, however, was in the context of a PCR data analysis, where confidence in the extrapolation probably does not reach the confidence level achievable with the data density possible with an isothermal reaction.

The visual impression of increasing response times for decreasing input target (Figure 4) is made quantitative on Figure 5D; the increased response time variance for lower input target levels is apparent as well. Greater response time variance at lower target levels is not due to ‘noisier’ data; the quality of the lines fit to the data is shown as correlation coefficient (R^2^) values on Figure 5F. The slope change with copy number is not so apparent on Figure 4, but is clearly seen on Figure 5F; while the intercepts (Figure 5G) showed no obvious relation to the input target level. RAM reaction rate decrease at low target concentrations has not been previously reported, and analysis of that rate decrease would have been difficult without the detailed data obtained as described.

A systematic change of slope for low target concentrations is consistent with the RAM reaction mechanism (5). In RAM reactions the amplification targets change dynamically. As the reaction proceeds, longer secondary templates undergo repeated transcription to produce double-stranded DNA products. Amplification reactions that are initiated from a single target molecule require the greatest progress through the substrates’ structure to accumulate sufficient double-stranded DNA product for fluorescence detection, and the difference in the substrate structure may underlie the change of slope with copy number. For example, a non-trivial probability of failure that is proportional to secondary template size would be a plausible mechanism for reaction rate reduction at lower initial target numbers. Experimental data (unpublished) suggests that the RAM reaction rate decrease at low input target number is probably not due to reagent exhaustion. The puzzling question, on first inspection of these data, was why there was so little intercept difference between target levels; and the slope dependence on target level was an obvious candidate for investigation.

The slope-intercept plots of Figure 6 were made to investigate the relation of target number, fitted line slope, and fitted line intercept in this experiment. When ordered by slope, the intercepts appeared to be roughly aligned within each input target level; and the slope dependence on input target is seen as a rightward shift (as panels are ordered on Figure 6) as copy number increases. The apparent intercept ordering by slope motivated a ‘second-order’ linear modeling, of intercept as a function of slope and copy number. The fitted lines plotted on Figure 6 were generated by one of those linear models. The endpoints of the lines were determined by the range of slope values; a copy-number of one was used to generate the line for the nominal 0.5 targets per reaction dilution set. Figure 6F shows that the slope-intercept data points, and their fitted lines, form separate diagonals on the combined plot. A gray line, perpendicular to the fitted lines, indicates a copy-number gradient. These models are statistical modeling of statistical abstractions; however, if copy-number and slope together can predict intercept, then slope and intercept may predict copy number.

Here the high frequency sampling data is used to investigate the behavior of RAM reactions at low copy number. Previous experiments with this system had shown diagonal patterns similar to the diagonals shown on Figure 6F. The target preparation and quantification followed by quantitative dilutions, described above, were undertaken to assess the formal possibility that the observed (and expected) RAM failures at high dilutions were failures to amplify two (or greater) target molecules per reaction. That situation might arise for example if, despite circularization, a sub-population of molecules were not competent to serve as targets. The results obtained showed that target copy number estimated from Poisson failure ratios was consistent with with target copy number estimated from spectrophotometic data and quantitative dilution; and the two dilutions where Poisson failure was observed are spatially separated on Figure 6F. It is therefore possible that reactions initiated from single target molecules are distinguished from reactions initiated from two or greater molecules on Figure 6F.

The applicability of these preliminary results within the domain of RAM methods remains to be determined. The results described, and similar unpublished results, have been obtained from a small subset of possible targets, primers, and amplification conditions. It is hoped, however, that these methods may suggest experimental and analytical approaches that will be useful for isothermal amplifcation in general. Statistical identification of a signal gradient that terminated in single target amplification would be generally useful for nucleic acids quantification. Recent analytical methods applied to real-time PCR (11, 12) should also be applicable to isothermal amplification, and those methods will likely benefit from the per-reaction efficiency measures obtained here.

## Methods

**ssDNA-enriched target preparation.** C-probe Cpr8FVWt1 (8) was circularized using CircLigase as recommended by the CircLigase supplier (Epicentre, Madison, Wisconsin). Following circularization, the reaction was split into aliquots for either exonuclease digestion of remaining linear C-probe by exposure to ExoIII (New England Biolabs (NEB) Ipswich, MA), or for mock exonuclease treatment (parallel preparation, but without exonuclease enzyme). Following exonuclease or mock-exonuclease treatment, both samples were recovered from Qiagen QIAquick nucleotide removal columns. Concentrations of single stranded circular C-probe molecules in the exonuclease-treated sample were estimated by absorbance measurement in a Nanodrop (NanoDrop products, Wilmington, DE) instrument. Absorbance readings were converted to a ssDNA concentration estimate using the molar extinction coefficient for the C-probe provided by the C-probe supplier (Gene Link, Inc.; Hawthorne, NY).

**Acrylamide gel electrophoresis** was done in 15% acrylamide-bis (19:1, Bio-Rad) in TBE; the 12 cm gel was run at 150 volts, 140 minutes, then stained in TBE running buffer with a 1:5000 dilution of Diamond(tm) Nucleic Acid Dye (Promega, Madison, WI) for 30 minutes. The gel image was captured on a Spectroline model TI-312E UV-transilluminator through with a Nikon Coolpix S52 digital camera and a Wratten No. 15 filter, 1 second exposure, f4.5, ISO 800.

**Quantitative dilutions** were prepared by a series of weighings on a Mettler AT201 analytical balance; all weighings were done in triplicate. Empty tubes were weighed, then diluent (10 mM TrisHCl, pH8 at 25°C; 1 mM EDTA) was added to tubes and a second series of weighings was done. Source solution was added to tubes followed by a third series of weighings.

**Real-time RAM** amplification reactions were performed by combination of equal volumes of 2x concentration enzyme-reaction buffer mix and of 2x concentration RAM primers plus target. Enzyme-buffer mix was 2x NEB isothermal amplification buffer, 200 uM each dNTP, 0.25x EvaGreen (Biotium Inc., Hayward, CA), and 0.26 units/ul ‘Warm Start’ Bst polymerase 2 (NEB). Primer-target mix was 5 uM Tris-HOAc pH 7.6 at 25°C, 0.1% Triton X-100 (), 1 nM fluorescein (BioRad), 1.75 uM forward primer Cpr8FVFwd61_22 (5’ACACTTCCAAACTCTCTCAATC), 1 uM Cpr8FVRvs84_19 (5’ACCTGTATTCCTCGCCTGT), plus the assay targets to the indicated levels. 10 ul of each 2x preparation were added to the wells of a 96-well microtiter plate for amplification. Real-time amplification reactions were performed in a BioRad (Hercules, CA) iCycler. The instrument was set to read continuously during 120 one-minute cycles.

**Real-time PCR** example reaction was performed on ∼950 ssDNA circularized targets described above,. Primers 0.75 uM each were Cpr8FVFwd70_20 (5’TTCACGCCTACACTTCCAAA), Cpr8FVRvs25_21 (5’AATACGAGAACACCCGATTGA).

OneTaq Hot Start polymerase (NEB), 0.05 units/ul in a 20 ul reaction); OneTaq standard buffer (3235, x); dNTPs (200 uM, each); Fluorescein (1 nM)). PCR parameters were 94°C, 20 seconds; 56°C, 30 seconds, and 68°C, 15 seconds. The ‘take-off-point’ and Second Derivative Maximum (SDM) points annotated on Figure 1D were obtained using the qpcR (14) package.

**Data analysis.** Data were extracted from the iCycler data file after using the iCycler software graphical user interface to deselect all filtering and to display all data points. The data was exported to a comma-separated values file, and imported into an R (13) data frame. Reactions were recorded as amplified or unamplified by determining signal change between the initial (baseline) data and terminal (plateau) data. Analysis of amplified reactions was done by numerical differentiation to find a maximum first derivative for the log(signal) vs time data. A (time, signal) point identified by the first derivative maximum was used as the midpoint of a 61 (time, data) values segment. Descriptive statistics for the segment (time of maximum first derivative, slope, intercept, goodness of fit, and variance were stored in a table for further analysis and figure production.

The statistical model that generated the lines on Figure 6 was generated by fitting the data from the three intermediate target levels (32, 8, and 2 nominal targets per reaction) to a model of the form

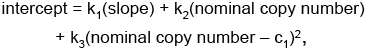

where k_1_, k_2_, k_3_, and c_1_ are numeric constants. Inclusion of the quadratic term for nominal copy number significantly increased the goodness of fit. The lines on the panels of Figure 6 were generated by computing the intercept for the maximum and minimum slope within each nominal copy number. Because the equation has the single factor k_1_ for the slope and the remaining terms constitute an added constant within each copy number, the generated values for the minimum and maximum slope determine the displayed fitted line.

## Acknowledgements

The author thanks Drs. Gail Radcliffe, James Smith, and Harry McCoy for critical reading of the manuscript.

